# Decoding multi-limb movements from low temporal resolution calcium imaging using deep learning

**DOI:** 10.1101/2023.11.30.569459

**Authors:** Seungbin Park, Megan Lipton, Maria C. Dadarlat

## Abstract

Two-photon imaging has been a critical tool for dissecting brain circuits and understanding brain function. However, relating slow two-photon calcium imaging data to fast behaviors has been challenging due to relatively low imaging sampling rates, thus limiting potential applications to neural prostheses. Here, we show that a recurrent encoder-decoder network with an output length longer than the input length can accurately decode limb trajectories of a running mouse from two-photon calcium imaging data. The encoder-decoder model could accurately decode information about all four limbs (contralateral and ipsilateral front and hind limbs) from calcium imaging data recorded in a single cortical hemisphere. Furthermore, neurons that were important for decoding were found to be well-tuned to both ipsilateral and contralateral limb movements, showing that artificial neural networks can be used to understand the function of the brain by identifying sub-networks of neurons that correlate with behaviors of interest.

## 1 INTRODUCTION

Decoding neural activity into behaviorally-relevant variables such as speech or movement is an essential step in the development of brain-machine interfaces (BMIs) and can be used to clarify the role of distinct brain areas in relation to behavior (Livezey and Glaser, 2021). In essence, “decoding” describes the use of an algorithm to translate between patterns of neural activity and behavior. To successfully achieve this goal, the algorithm must identify a population of neurons that contains information about the behavior of interest. Then, examining the response properties of neurons that are important to the decoding algorithm should illuminate the way in which information is encoded within the brain.

Behavior emerges from the interaction of neural circuits within and across brain areas (Peron et al., 2015); therefore, recording from large neural populations is essential to improving neural decoding accuracy (Pandarinath, O’Shea, et al., 2018). For example, two-photon (2p) calcium imaging enables recording the activity of thousands of neurons with a single-cell resolution in genetically-defined populations (Hofer et al., 2011; Peron et al., 2015; Stringer et al., 2019; Weisenburger et al., 2019). As a result, 2p calcium imaging has advanced our understanding of circuit-based neural processing underlying, for example, movement (Huber et al., 2012; Li et al., 2019; Omlor et al., 2019; Zhu et al., 2022), proprioception (Alonso, Scheer, Palacio-Manzano, et al., 2023; Mamiya et al., 2018), touch (Peron et al., 2015), and vision (de Vries et al., 2020; Stringer et al., 2019). Given these advantages, next-generation optical BMIs have looked towards 2p calcium imaging as means for neural recording (Clancy et al., 2014; O’Shea et al., 2017; Trautmann et al., 2021). However, decoding 2p calcium imaging recordings for use in real-time applications has traditionally been challenging due to the slow sampling rate of the signal (Siegle et al., 2021) and the indirect and non-linear relationship between underlying neural activity and the slow fluorescent signal (Chen et al., 2013). In consequence, the major focus in processing 2p calcium imaging data has been to develop methods to infer the underlying low-dimensional dynamics of the neural population to improve decoding of movements from neural activity patterns (Schneider et al., 2023; Zhu et al., 2022). However, direct decoding of neural activity from 2p imaging data is possible using artificial neural networks (ANNs) (Li et al., 2019; Liu et al., 2022).

Seminal work in the field of BMIs traditionally relied on linear methods including the Kalman and Wiener filters for decoding movement intention from the spiking activity of neurons (Carmena et al., 2003; Gilja et al., 2012; Orsborn et al., 2014; Yu et al., 2009). Recently, however, the use of machine learning and deep learning has become popular given the non-linearity in the task and the development of deep learning tools (Livezey and Glaser, 2021; Makin et al., 2020; Pandarinath, O’Shea, et al., 2018). Deep learning has also been successfully applied to decode the movement of animals from 2p imaging data. For example, a convolutional neural network (CNN) can accurately decode (classify) forelimb reach endpoints from 2p imaging (Li et al., 2019). Various types of ANNs can decode forelimb movement during a simple lever-press task (Liu et al., 2022). Furthermore, low-dimensional neural population dynamics inferred from 2p imaging using a deep learning framework can be linearly related to forelimb reach trajectories (Schneider et al., 2023; Zhu et al., 2022). However, the behavioral variables that have been decoded from 2p imaging to date are limited: either discrete endpoints or repetitive reaches of single limbs rather than continuous, dynamic, and naturalistic multilimb movements.

Despite the common simplification that sensorimotor cortices primarily encode and control the contralateral movement of the body, these brain areas additionally contain information about ipsilateral limb movements (Ames and Churchland, 2019a; Gale et al., 2021; Halley et al., 2020) that emerges from trans-callosal projections from the primary somatosensory cortex (S1) in the contralateral hemisphere (Naskar et al., 2021). Ideally, such multi-limb information could be used to control a complex, naturalistic multi-limb BMI. Therefore, in this present study, we set out to validate the theory that a deep learning approach could be used to decode the continuous multi-limb trajectories of running mice from neural recordings made with 2p calcium imaging over S1 and primary motor cortex (M1). Furthermore, we aimed to link the decoder function to the properties of neural tuning within mouse S1/M1, effectively using the decoding to infer properties of neural coding within these cortical areas.

To achieve these goals, we used a recurrent encoder-decoder model with Long Short-Term Memory (LSTM), in which the output sequence length of the decoder was longer than the input sequence length to the encoder (Makin et al., 2020; Sutskever et al., 2014). This method overcame the low sampling rate of 2p calcium imaging relative to behavior, and it was able to predict the movements of all four limbs from neural recordings in a single cortical hemisphere. Moreover, we improved the interpretability of the deep learning model by extracting the importance of neurons for decoding from the trained models (Makin et al., 2020) and then comparing neural tuning between the most and least important neurons. The results show that the small fraction of highly important neurons selected by the encoder-decoder model yields optimal decoding performance and is well-tuned to limb movements, validating decoding by the encoder-decoder models was not by chance. Our work demonstrates the feasibility of using deep learning methods to identify and characterize populations of neurons that encode behaviorally-relevant variables. This approach will be critical in the future implementation of neural decoding for next-generation optical BMIs that will improve the lives of patients suffering from neurological injury and disease.

## 2 RESULTS

### Overview of the dataset and the deep learning framework

We used 2p calcium imaging to record neural activity from S1 and M1 of TRE-GCaMP6s x CaMKII-tTA mice while the animals were head-fixed but freely running on an air-lifted ball (Figure 1A). Animal running behavior was recorded using bilateral cameras capturing frames at 30 Hz, and neural activity was recorded simultaneously with a Neurolabware mesoscope at 7.8 Hz. The experiment was conducted on three subjects (Mouse A, Mouse B, and Mouse C). The total length of the recordings was 20 minutes 24 seconds (36731 behavioral video frames and 9419 neural imaging frames) for Mouse A, 20 minutes 12 seconds (36374 behavioral video frames and 9315 neural imaging frames) for Mouse B, and 20 minutes 9 seconds (36297 behavioral video frames and 9305 neural imaging frames). Figure 1B shows the imaging planes of each mouse across sensorimotor cortical areas. Recordings were targeted to stereotaxically defined S1, although the recording from Mouse C also covered a portion of M1. In each mouse, single neurons could easily be identified from the imaging frames (Figure 1C). The identified number of neurons was 3327, 2978, and 3314 for Mouse A, Mouse B, and Mouse C, respectively.

**Figure 1.**
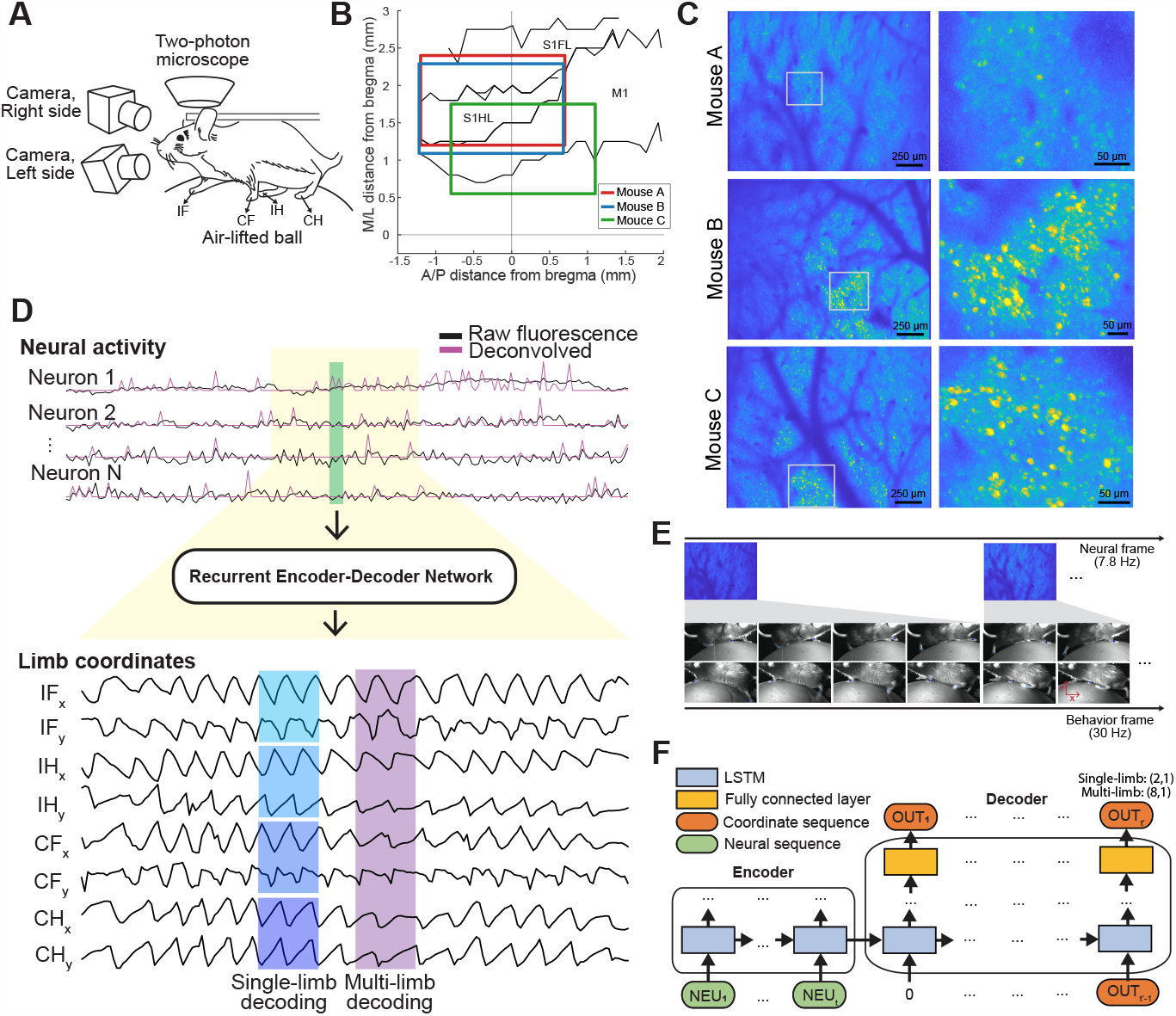
The 2p imaging dataset with behavior and the overall framework. (A) Recording setup. Mice ran on an air-lifted ball while neural activity was recorded with a 2p mesoscope and behavior was recorded with cameras on each side of the body. IF: ipsilateral front limb; IH: ipsilateral hind limb; CF: contralateral front limb; CH: contralateral hind limb. (B) Imaging planes for 2p imaging on the cortical map of a mouse. S1FL: the primary somatosensory cortex area for a front limb; S1HL: the primary somatosensory cortex area for a hind limb; M1: the primary motor cortex. (C) Maximum intensity projections of recorded neurons during 2p imaging. (D) The overall framework showing input (neural activity from 2p imaging), the encoder-decoder model, and outputs (limb coordinates). Neural activity with a yellow background corresponds to the limb coordinates plotted in the figure. The green box indicates the input to the encoder-decoder model. Blue and purple boxes in limb coordinates show outputs from the encoder-decoder model — while these sequence lengths match the sequence length of the neural activity, the timing of the input and the output does not temporally match for visualization purposes. (E) Limitation of low sampling rate of 2p calcium imaging. Because the neural signal has a much lower sampling rate than the behavior, one neural frame matches multiple (e.g. four) behavior frames. (F) The schematic diagram of the encoder-decoder model — the recurrent encoder-decoder network. The length of the input neural sequence (t) is smaller than the length of the output coordinate sequence (t’).

To address the challenges related to decoding 2p calcium imaging recordings, we designed a decoding approach that uses deep learning to decode limb movements from neural activity (Figure 1D). The encoder-decoder model takes deconvolved neural activity as inputs and outputs the x- and y-coordinates of one (single-limb decoding) or four limbs (multi-limb decoding): ipsilateral (right) front (IF*x* and IF*y*), ipsilateral hind (IH*x* and IH*y*), contralateral (left) front (CF*x* and CF*y*), and contralateral hind limbs (CH*x* and CH*y*). The large discrepancy between the low 2p imaging sampling rate and the high behavioral video sampling rate represents the challenge we aim to address in this study (Figure 1E): only one neural imaging frame corresponds to multiple behavior video frames (e.g. four frames). Considering the discrepancy between the sequence length of neural signals (input) and behavior variables (output), we designed a recurrent encoder-decoder network (Figure 1F). This encoder-decoder model was inspired by a sequence-to-sequence learning model for machine translation (Sutskever et al., 2014) that has been previously applied for decoding speech from neural activity (Makin et al., 2020). The encoder-decoder model has two parts: 1) the encoder and 2) the decoder, both with LSTM, which allows different input and output lengths. Single-limb decoding models for each limb and multi-limb decoding models for all limbs were trained for Mouse A, Mouse B, and Mouse C, respectively (Supplemental Table 1). Decoding performance was evaluated qualitatively and quantitatively on separate test data (20%) that was not included in the training set.

### Single-limb decoding of contralateral and ipsilateral limb trajectories

First, we tested whether the designed encoder-decoder model can decode limb trajectories from fluorescence traces extracted from 2p imaging recordings. Particularly, we focused on whether both contralateral and ipsilateral limbs can be decoded from a unilateral cortical hemisphere recording. Using encoder-decoder models trained to decode the movement of single limbs (single-limb decoding), we found that the single-limb decoding models accurately decoded both x- and y-coordinates of limb position over time for each of the three mice. As a metric for performance, root mean squared error between the processed ground truth and the processed predictions was calculated (RMSEp, see Methods). For qualitative evaluation, partial sequences (200 frames) of ground truth and predictions with the best average RMSEp over all coordinates were plotted for each mouse (Figure 2A). Visualization of the sequences showed that there was variability not only in decoding accuracy but also in the ground truth of different sequences across limbs and mice. For example, while the characteristics of signals in Mouse A seem to be similar across limbs with regular 4 Hz oscillations, there was more irregularity in Mouse B and Mouse C.

**Figure 2.**
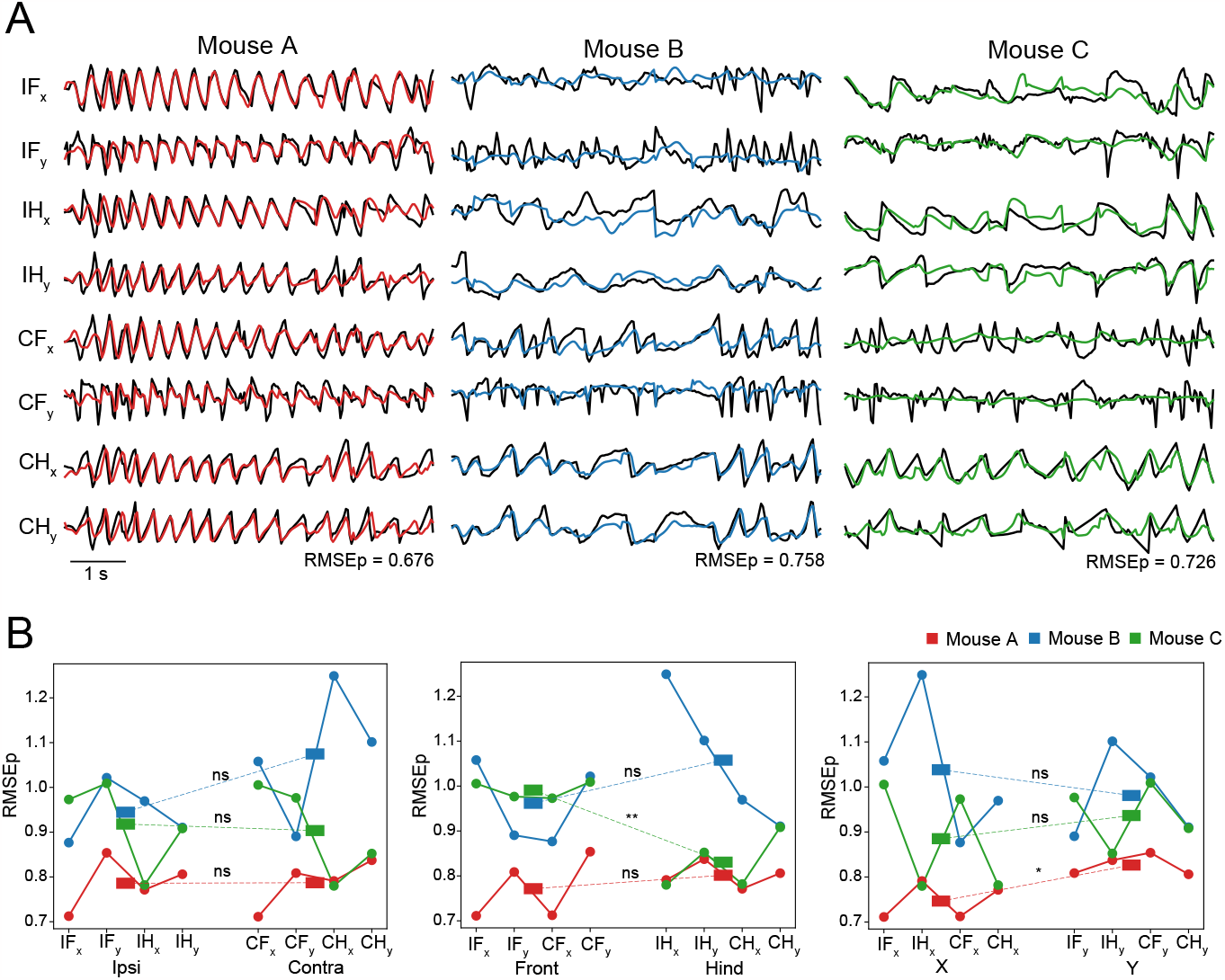
Single-limb decoding of contralateral vs. ipsilateral limbs. **(A)** Sample traces showing ground truth (black) and predictions (colored) by the encoder-decoder models for contralateral and ipsilateral movement in mouse A, B, and C. **(B)** Limb prediction RMSEp across all mice and all limbs, grouped by ipsilateral vs. contralateral (left), front vs. hind limbs (middle), and x-coordinates vs. y-coordinates of limb position (right). Stars indicate significance: ns *p >* 0.05, * *p <* 0.05, ** *p <* 0.01; t-test.

The evaluation of performance was quantitatively represented by RMSEp (Figure 2B). Corresponding to the qualitative evaluation, Mouse A showed the best performance with the lowest average RMSEp over all coordinates. We found no net significant difference between decoding error for contralateral vs. ipsilateral limbs (the left and right limbs of the mouse, respectively, given window placement on the right cortical hemisphere) (All mice: *p* = 0.479; Mouse A: *p* = 0.979; Mouse B: *p* = 0.157; Mouse C: *p* = 0.849; t-test), between front vs. hind limbs (All mice: *p* = 0.835; Mouse A: *p* = 0.465; Mouse B: *p* = 0.318; t-test), or between x- and y-coordinates of limb position (All mice: *p* = 0.654; Mouse B: *p* = 0.563; Mouse C: *p* = 0.489; t-test). There was only a significant difference in front vs. hind limbs of Mouse C (*p* = 0.003; t-test) and x-vs. y-coordinates of limb position of Mouse A (*p* = 0.014; t-test). This result demonstrates that the encoder-decoder model could decode all contralateral and ipsilateral front and hind limb trajectories.

### Simultaneous multi-limb decoding of multiple limb trajectories

The encoder-decoder model’s success at accurately decoding each of the animal’s four limbs shows that sufficient information about both ipsilateral and contralateral limbs is contained within the neural population from a single cortical hemisphere. Next, we trained the encoder-decoder model to output the x- and y-coordinates of all four limbs simultaneously (multi-limb decoding), where it has previously decoded only a single limb at a time (single-limb decoding). The same metric of evaluation for decoding accuracy used for single-limb decoding, RMSEp, was computed for the multi-limb decoding model (Supplemental Figure 1). We found no significant difference in decoding accuracy between the single-limb decoding RMSEp and the multi-limb decoding RMSEp (*p >* 0.05, Wilcoxon rank-sum test; Figure 3). These results are surprising, given that decoding more limbs would theoretically require using fewer neurons to decode each limb, thus reducing decoding accuracy (Schneider et al., 2023; Zhu et al., 2022). We predicted that similar decoding performance between the single-limb and multi-limb decoding may occur if the encoder-decoder models rely on only a fraction of the neural population that is essential for decoding.

**Figure 3.**
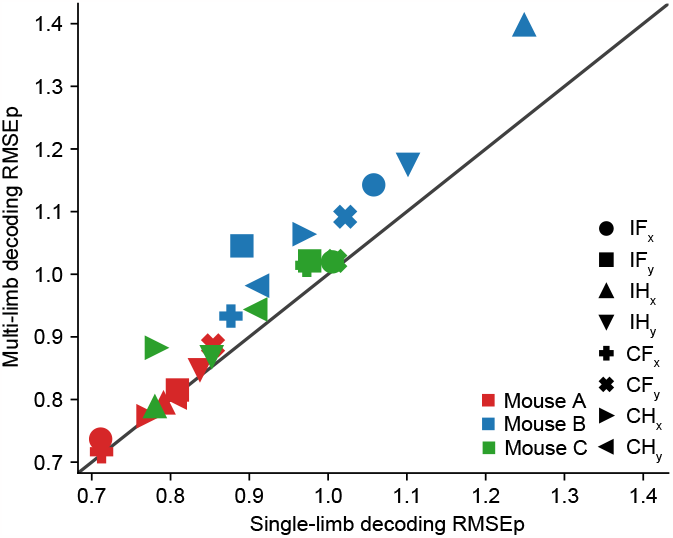
Summary RMSEp for single-limb decoding vs. multi-limb decoding

### The encoder-decoder model relies on only a fraction of neurons within the population for accurate decoding

To determine how many and which neurons are used by the encoder-decoder models, we calculated the relative importance of neurons for decoding in the multi-limb decoding models using the gradients of the loss function with respect to each neuron (see Methods; Makin et al., 2020; Simonyan et al., 2014). The distributions of importance from multi-limb and single-limb decoding models are shown and compared in Figure 4A, and the difference was significant (multi-limb vs. single-limb combined for each animal, *p <* 0.0001, Kolmogorov-Smirnov test). The importance from single-limb decoding models was more skewed leftwards than the multi-limb result. We found that the distribution of neural importance values was skewed leftwards even in the multi-limb decoding result, centered between 0.1 to 0.2, with a fat tail towards the maximum value (normalized to one within each animal; Figure 4A). This asymmetry suggests that, even in the multi-limb decoding model, only a fraction of neurons are relied upon for decoding limb trajectories.

**Figure 4.**
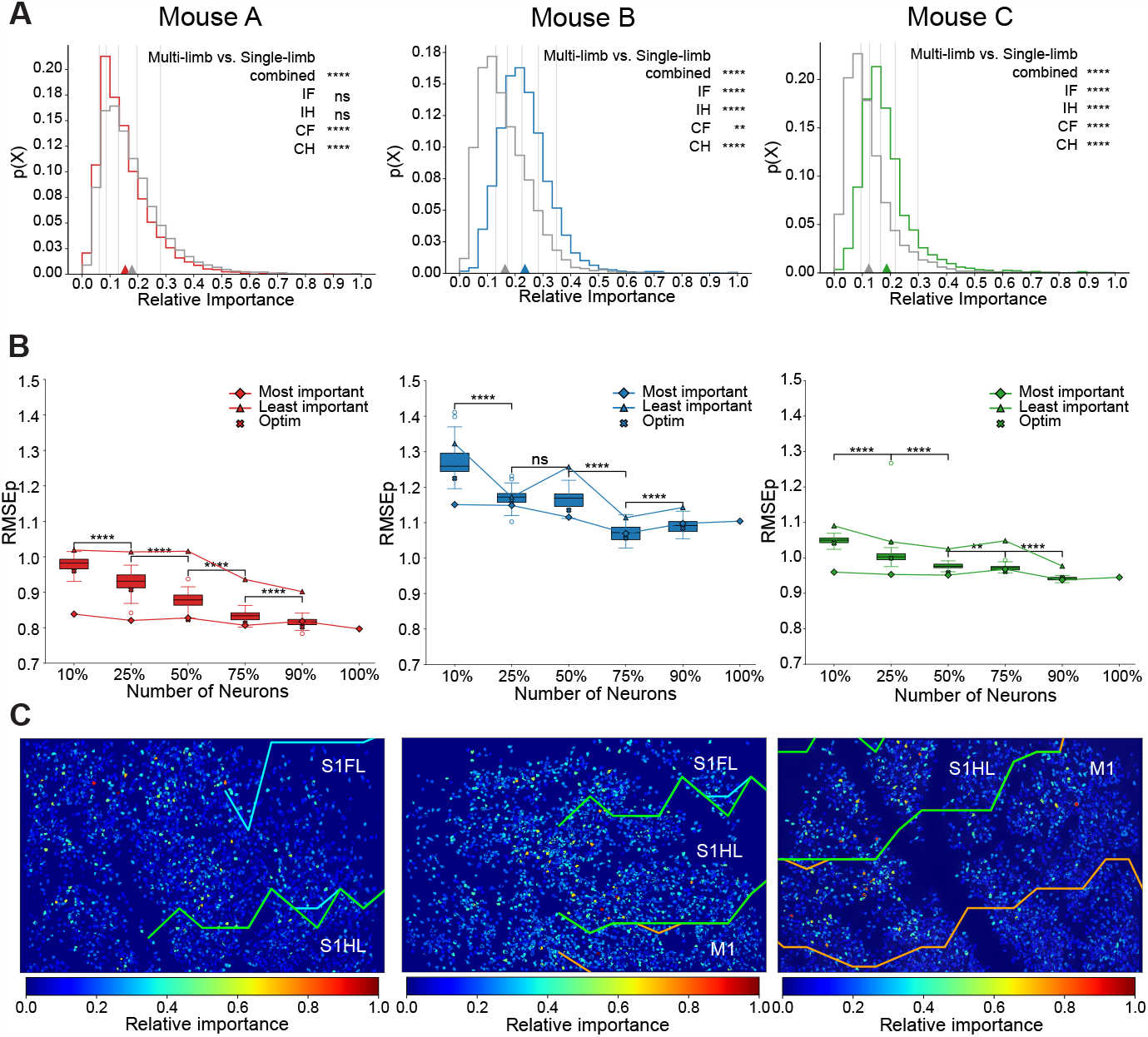
Relative importance of neurons distributed over the imaging planes. **A)** The distribution of normalized neural importance of multi-limb decoding models (colored) and single-limb decoding models (grey), shown separately for each animal. Vertical bars show the top percentiles (10%, 25%, 50%, 75%, and 90% from right to left). Stars indicate the significance of the difference between multi-limb (colored) and single-limb (grey) models of each limb (IF, IH, CF, and CH) or all-limbs combined (combined): ns *p >* 0.05, ** *p <* 0.01, **** *p <* 0.0001; Kolmogorov-Smirnov test. **B)** Decoding accuracy using subsets of neurons, broken into the most important neurons (line with diamonds), the least important neurons (line with triangles), and randomly-sampled neurons (box plots). One of the randomly sampled sets was used for optimization of the encoder-decoder models, i.e. hyperparameter tuning (Optim). Relationship between the fraction of neurons used in decoding and RMSEp shown separately for each Mouse A, B, and C (colors as in (A)). Stars indicate significance: ns *p >* 0.05, ** *p <* 0.01, **** *p <* 0.0001; t-test. **C)** Heat maps of the relative importance of neurons plotted over the imaging planes. S1FL: the forelimb area of S1; S1HL: the hindlimb area of S1; M1:

To better understand how the encoder-decoder model performance depends on the population of neurons that is used for decoding, we re-trained the multi-limb decoding models on data consisting of different subsets of neurons. One of the randomly sampled subsets was used for hyperparameter tuning. Using this fixed set of hyperparameters, we trained the encoder-decoder models on the top 10%, 25%, 50%, 75%, and 90% of the most and the least important neurons. As expected, we found that going from 10% to 90% of neurons often yielded only a small improvement in decoding accuracy (Figure 4B). Next, using the same set of hyperparameters, the encoder-decoder model was trained on 50 additional randomly-selected subsets of the population to compare decoding with important neurons to decoding with a randomly-sampled population. As expected, the RMSEp of these trained encoder-decoder models is proportional to the number of neurons being used, with errors significantly decreasing as additional neurons are added (aligning with Zhu et al., 2022 and Schneider et al., 2023). Furthermore, the highly important neurons significantly out-perform randomly-selected neurons at 10% and 25% (although the latter only holds true for mice A and C), while the low-important neurons show the worst performance. Given that the encoder-decoder model trained with highly-important neurons can achieve high decoding accuracy using just 10% of neurons, it is likely that randomly-selected subsets of, e.g., 50% of neurons may overlap with enough of the “important” neuron population to allow both models to perform well.

Since our imaging window spanned multiple cortical areas (although predominately S1 and M1), we decided to test whether the most important neurons were located within a specific cortical area or whether they were distributed across the imaging field. To address this question, we plotted the spatial distribution of important neurons, color-coding each neuron by its relative importance (Figure 4C). These plots show that highly important neurons are sparsely distributed over the cortex area. To confirm the sparse distribution of the space, the imaging planes were divided into grids with a length of 50 pixels, and the relative neural importance was averaged in each grid (Supplemental Figure 2). The grids show that the spatially averaged values are in a similar range, confirming that neurons important for decoding are sparsely and evenly distributed over the cortex.

### High-importance neurons are well-tuned and correlated with limb movements

To better understand why some neurons are more relied upon for neural decoding of limb trajectories, we compared the tuning properties of the top ten percent of the most and least informative neurons, choosing the value of ten percent given the high decoding accuracy with that fraction of neurons (Figure 4B). A visual inspection of the neural activity for the most and least important neurons during limb movements suggests differences in neural activity patterns between the two groups (Figure 5A): the activity of important neurons is strongly correlated with animal behavior, with high-activity periods during motion and low activity while the animal is still. In contrast, the least important neurons are active throughout the entire recording session or have low-activity periods that are uncorrelated with animal behavior. We quantified these observations by calculating the total spiking activity of each neuron (Figure 5B) and the correlation between each neuron and the movement of each limb (separately for x- and y-coordinates; Figure 5C). Supporting our qualitative observations, important neurons had statistically lower overall activity levels (likely from having low activity while the mice were still) and were statistically more correlated with limb movements than were the least important neurons (Figure 5B,C). Thus, the encoder-decoder model uses information from neurons whose activity is best related to limb movements.

**Figure 5.**
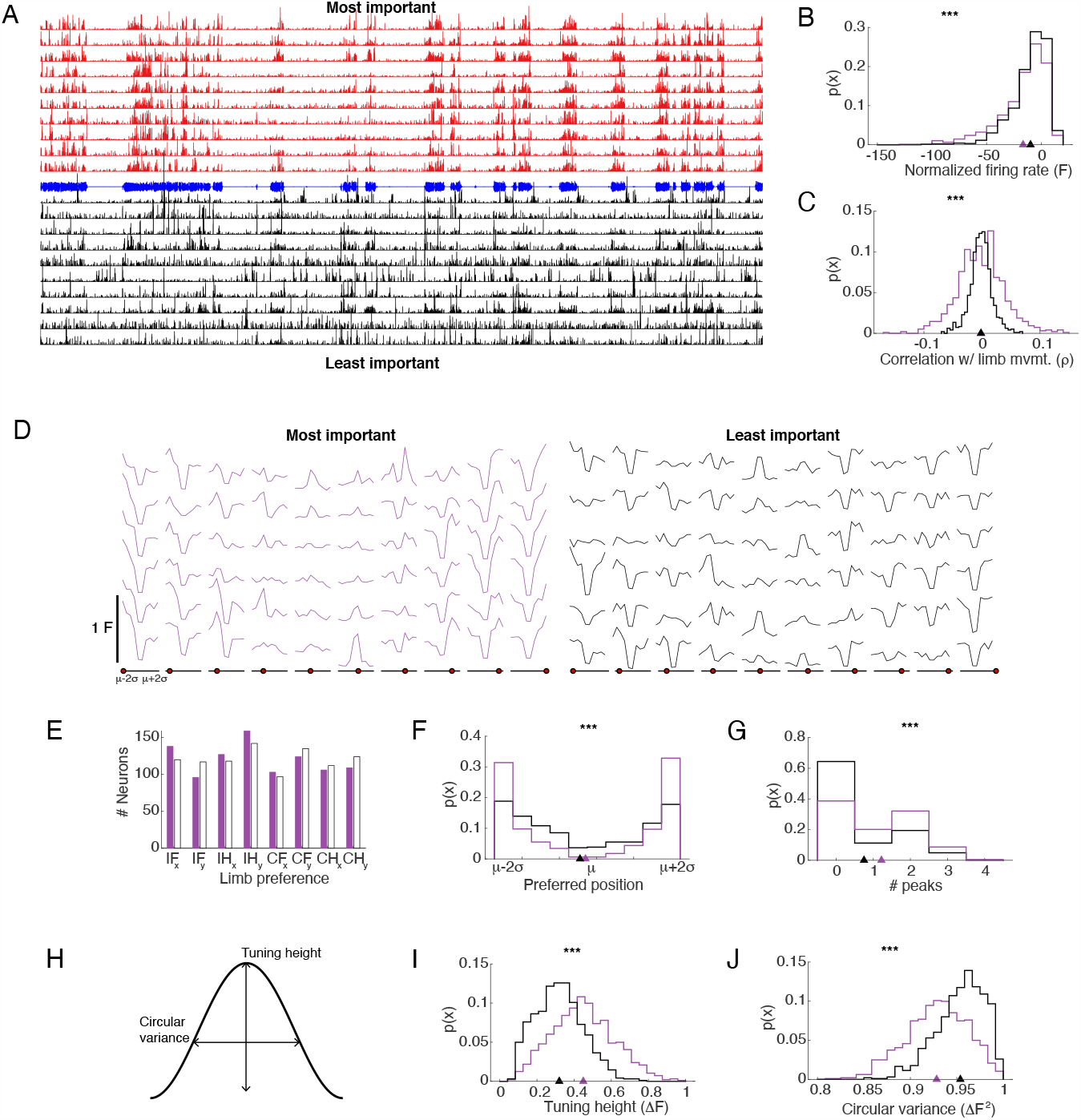
Tuning properties of important neurons. **A**. Sample normalized and deconvolved neural response traces, shown for the ten most (red, top) and least (black, bottom) informative neurons from Mouse A. Neural activity traces can be compared to limb movement traces, shown in blue (contralateral hindlimb movement of Mouse A along the x dimension). **B**. Distribution of the average neural firing rates for the most (purple) and least (black) important neurons (*** *p ≤* 0.00001, Wilcoxon rank sum test). Data was consolidated across all mice. **C**. Correlation between neural activity and limb movements (see Methods: neural correlations). Colors as in (B) and data was consolidated across all mice. *** *p ≤* 0.00001, Wilcoxon rank sum test. **D**. Sample tuning curves to limb position. Each column contains neurons with a different preferred location, going from *µ −* 2*σ* at far left to *µ* + 2*σ* at far right. The preferred location in each column is indicated by the position of the red circle along the line showing position from *µ −* 2*σ* to *µ* + 2*σ*. Purple shows tuning curves for a subset of the top ten percent most important neurons. Black shows tuning curves for a subset of the least important neurons. Tuning curves are shown for each neuron’s preferred limb. **E**. Distribution of preferred limbs for the most (purple) and least (black) important neurons, consolidated across all mice. **F**. Distribution of preferred limb positions for each neuron’s preferred limb for the most (purple) and least (black) important neurons, consolidated across all mice. *** *p ≤* 0.00001, Kolmogorov–Smirnov test. **G**. Distribution of the number of significant peaks in the tuning curves (where two peaks indicates bimodal and zero peaks indicates untuned) for the most (purple) and least (black) important neurons, consolidated across all mice. *** *p ≤* 0.00001, Kolmogorov–Smirnov test. **H**. Schematic depicting two measures of neural tuning to limb position: tuning curve height and circular variance, the latter of which measures the spread of the neural responses across the possible positions. **I**. Distribution of the tuning curve heights, calculated as the difference between the maximum and minimum response. Colors as in (B) and data was consolidated across all mice. *** *p ≤* 0.00001, Wilcoxon rank sum test. **J**. Distribution of the tuning curve circular variance, which is a measure of the width and spread of the tuning curve. Colors as in (B) and data was consolidated across all mice. *** *p ≤* 0.00001, Wilcoxon rank sum test.

To further analyze the relationship between neural activity and behavior, we calculated tuning curves for each neuron as a function of limb position (Figure 5D). We then compared the tuning properties of the top and bottom ten percent of neurons, sorted in order of normalized importance. First, we found that both the most and least important neurons were nearly uniformly distributed in terms of the limb to which they best responded (Figure 5E). Next, both types of neurons tended to be tuned towards large deviations from the mean position, with only a smaller fraction tuned to the center (Figure 5F). The asymmetry in preferred limb positions (the x or y position to which a neuron had the largest response) was significantly more pronounced for the most important neurons over the least important neurons. In general, the distribution’s skew towards the extremities suggest that neural activity in S1/M1 is encoding limb position as a function of the change from baseline position. Furthermore, many of the neural tuning curves tended to be bi-modal (Figure 5D,G), with peaks for movements both anterior and posterior to the baseline. Bimodal tuning was more common for the most important versus the least important neurons (Figure 5G). These results align with prior work examining the representation of proprioception within mouse S1/M1: neurons tend to respond both to movements towards and away from the body (Alonso, Scheer, Palacio-manzano, et al., 2023). Finally, we measured two important properties of the neural tuning curves to position: tuning height and the width or spread of the tuning curve, the latter of which we measured using circular variance (Figure 5H). Circular variance reflects the width or spread of the curve, so smaller circular variances reflect narrower tuning and large circular variances reflect wider tuning. As predicted, we found that highly-important neurons tended to have tuning curves that are narrower with higher peaks than the least important neurons. A comparison of the distribution of tuning heights showed that most important neurons have larger tuning height values compared to least important neurons (Figure 5I). Additionally, comparison of the distribution of circular variance showed that most important neurons tended to have smaller circular variance values compared to least important neurons (Figure 5J). Therefore, the encoder-decoder model identified neurons that are well-tuned to the behavioral task of interest: neurons that are strongly correlated with behavior, have low unrelated activity levels, and have strong and selective responses to limb position.

### Limb movements with higher frequency content can be accurately decoded

To examine the ability of the encoder-decoder model to decode behavioral data in the form of limb movement in finer detail, we used a Fourier transform to compute the frequency distribution of individual limb trajectories (separately for x and y position) and of the predictions of limb trajectories made by the multi-limb decoding model (Figure 6A). We found that, while both the true and predicted limb movements had the highest power at frequencies around or below the Nyquist frequency of the 2p imaging dataset (ImagingNyquist = 3.9 Hz), there was significant frequency content above the imaging Nyquist in both the behavioral data and in the model predictions. Clearly, given the Nyquist theorem (Nyquist, 1928), this information is not available from the time-varying activity of single neurons. Instead, our ability to predict high-frequency movement suggests that information about movements is encoded in a distributed fashion across the population of neurons. If that were true, the encoder-decoder model could decode limb trajectories from the identity of neurons that were active in a given imaging frame rather than from the time-varying activity of individual neurons. In consequence, by using data recorded across the population of neurons, the encoder-decoder model can predict higher-frequency information than is nominally available in the neural recordings.

**Figure 6.**
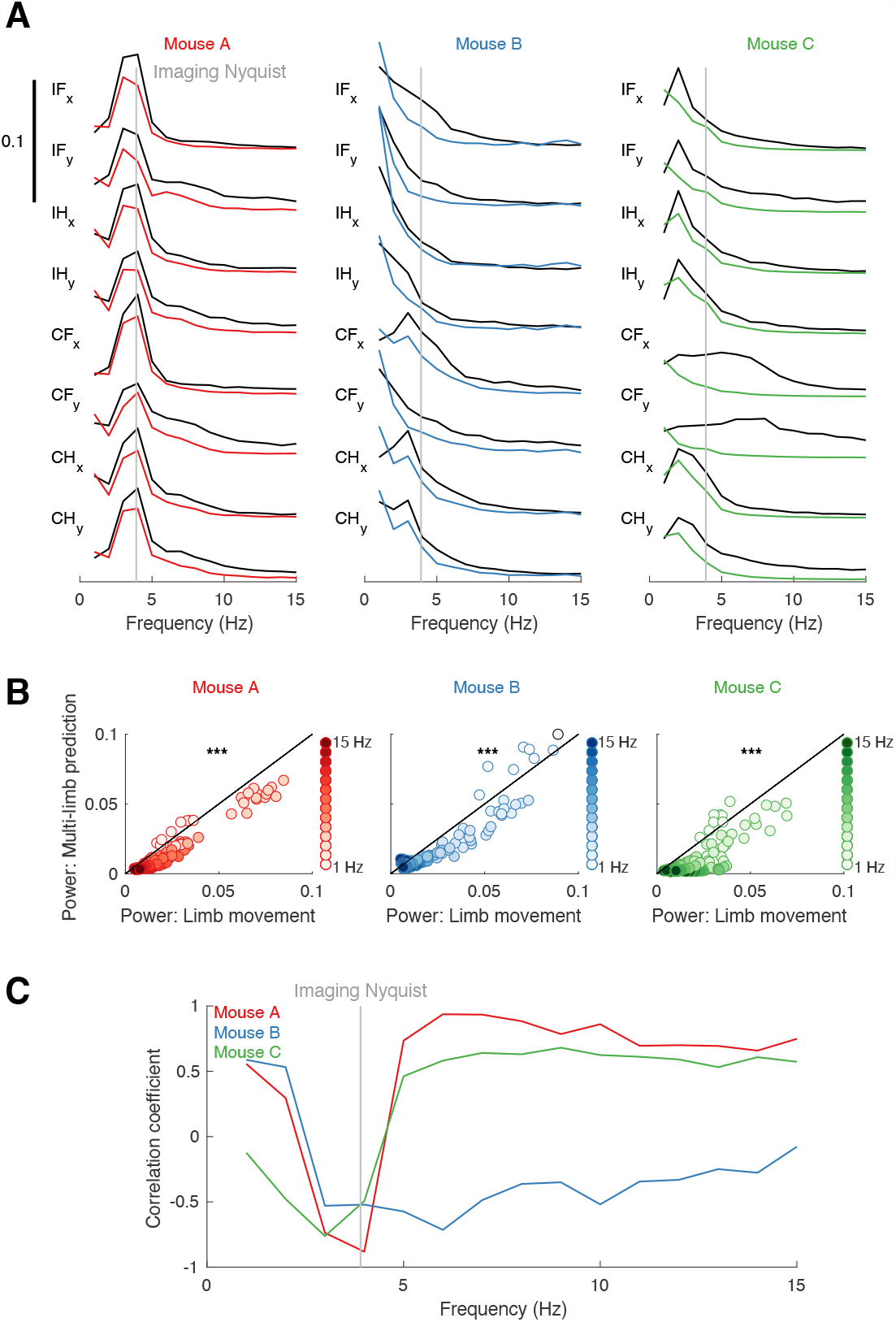
Power spectrum of frequency content of behavioral movements and predictions. **A**. Power spectrum of limb movements (as labeled) and multi-limb decoding model predictions for each of the three subjects. Black lines indicate ground truth (true limb movements) while colored lines indicate multi-limb decoding model predictions. Vertical gray line shows the Nyquist limit for the 2-photon recordings. **B**. Direct comparison of the power at each frequency between limb movements and model predictions, separated by subject. These plots include data from all four limbs and is identical to that presented in (A). *** *p ≤* 0.00001, Wilcoxon rank sum test. **C**. Correlation between the power at each frequency and multi-limb decoding RMSEp across limbs. Gray vertical line indicates the Nyquist frequency for the 2-photon recordings.

Despite the encoder-decoder model’s ability to predict high-frequency information, the predictions it makes have lower power across the frequency spectrum than the true behavioral data (Figure 6B). Only at the lowest frequency (1 Hz) does the encoder-decoder model have higher power than the true behavior, which shows that it often defaults to estimating the mean trajectory (as can be seen in Figure 2A). Finally, to identify the frequencies at which the encoder-decoder model makes the most accurate predictions, we correlated the power in each frequency band with decoding error at that limb (Figure 6C). In this analysis, a negative correlation indicates lower errors at higher levels of power at each frequency. We found a strong negative correlation around 3-4 Hz in all three mice, which we found was approximately the frequency of leg swings during running in the mice (data not shown). Although movement speed varied across mice and across time, this range of frequencies was common across all limb trajectories (see spectral decomposition of limb movements in Figure 6A). Therefore, it seems that the encoder-decoder model was learning the global structure of movements during locomotion as well as finer details of limb movements.

## 3 DISCUSSION

Although 2p calcium imaging enables stable recording of large populations of neurons at single-cell resolution, relating 2p imaging data to motor activity has been challenging due to the low imaging sampling rate relative to movement and due to the non-linear relationship between: 1) the underlying neural activity, 2) the 2p imaging signals, and 3) sensorimotor behavior. Here we demonstrated that a deep learning-based approach can successfully decode continuous, multi-limb trajectories of running mice from 2p calcium imaging recordings in S1/M1. Notably, ipsilateral and contralateral limb movements were decoded with similar accuracy (Figure 2), and there was no loss in accuracy when simultaneously decoding the movement of all four limbs compared to decoding the movement of single limbs (Figure 3).

### Comparison to prior work

Using deep learning to decode movements directly from 2p imaging data is a relatively new concept with few prior studies (Li et al., 2019; Liu et al., 2022). One such study used pre-trained Resnet18 to predict reach endpoints, using the raw 2p imaging frames as inputs to the model (Li et al., 2019). Using the raw 2p imaging frames can be beneficial for real-time applications and exploit the spatial information that is lost when extracting fluorescence traces from isolated regions of interest (i.e., identified neurons). However, CNNs ignore the temporal information embedded in time-varying continuous neural signals that may include critical information for movement. Furthermore, the classification of limb-reaching endpoints in clusters has limited applications. A second study applied multiple deep learning architectures in different decoding schemes (concurrent, time-delay, and spatiotemporal information) to decode forelimb movement during a simple lever-press task (Liu et al., 2022). In concurrent and time-delay schemes, they show ANN and LSTM perform better than other machine learning algorithms. Compared to using a single LSTM network, our approach uses two separate LSTM networks to account for the different input and output lengths, thereby overcoming the low sampling rate problem. Next, the authors attempted to utilize the spatial information by using a hybrid spatiotemporal network with ANN and CNN in which the spatial relationship or connectivity between neurons is provided as reconstructed images of neural activity and connectivity. However, this hybrid approach neglects the temporal information in that it uses ANN, not LSTM. An alternative to these approaches is to improve the 2p imaging neural signal by first inferring low-dimensional neural population dynamics that subsequently improve behavioral decoding accuracy (Zhu et al., 2022). This was achieved by applying a variational autoencoder with recurrent neural networks (RNNs) to elicit neural population dynamics from 2p calcium imaging recordings. In contrast, our study directly applies the encoder-decoder with RNNs to predict movement from the neural signals, which may lead to faster real-time implementation. Notably, all of these studies aimed to decode endpoints or movements of a single limb in simple reaching tasks. Thus, a major advantage of our current work is simultaneously decoding movement of all four limbs during complex, natural running from a single cortical hemisphere.

### Explainable encoder-decoder network: using artificial neural networks to understand neural circuit function

Instead of treating the encoder-decoder network as solely a means to an end (i.e., to translate between neural activity and intended movements in BMIs), it can be viewed as a means by which to examine neural encoding of behavior within the brain. Towards this aim, we calculated the dependence of the neural network on each identified neuron (neural importance in Figure 4) and then analyzed the response properties of important neurons (Figure 5). It is of note that the neurons identified as “important” by the trained encoder-decoder model share properties reported by other studies to be characteristic of neural coding of movement, including: 1) sparse distribution over the cortical areas (Peters et al., 2014), 2) a small fraction of the most important neurons are relied upon for decoding (Barth and Poulet, 2012; Brecht et al., 2004; Olshausen and Field, 2004; Takiyama, 2015), 3) bimodal tuning curves (Alonso, Scheer, Palacio-manzano, et al., 2023), and 4) narrow tuning curves (Meier et al., 2020; Montemurro and Panzeri, 2006).

Our results thus support known theories of sensorimotor coding, including “sparse coding” (Barth and Poulet, 2012; Brecht et al., 2004; Olshausen and Field, 2004; Takiyama, 2015), which states that only a small number of neurons are extensively activated for various sensory or motor processing. Furthermore, the bimodal tuning curves suggest that neurons may not be encoding specific limb positions in space so much as limb position relative to the body. A similar notion has been shown for neural coding of proprioception, where the direction and amplitude of a movement vector is encoded in S1/M1 rather than a specific start and end (Alonso, Scheer, Palacio-manzano, et al., 2023). Lastly, tuning sharpness or selectivity, which emerges from recurrent cortical connections (Somers et al., 1995), is related to neural encoding of information (Meier et al., 2020; Montemurro and Panzeri, 2006). Intuitively: if a neuron responds strongly to a small range of limb positions, we can be certain that the limb is in one of those positions when that neuron is active. In contrast, if the neuron responds strongly to a broad range of limb positions, then that neuron’s activity is less informative about the time-varying position of the limb. The encoder-decoder model thus supports the notion that narrow tuning curves contribute to more efficient encoding of sensorimotor information in the brain. Finally, although we do not test this hypothesis here, the model’s ability to extract higher-frequency movements than should be possible according to the Nyquist theorem strongly suggests that movement sequences are encoded in the joint or sequenced activation of a group of neurons rather than the time-varying activity of neurons.

The close match between our analysis of encoder-decoder function and past experimental work validates that the trained decoding scheme of encoder-decoder models was not random or simply extracting certain patterns in limb trajectories, but that the model indeed exploited the neural activity recorded by 2p calcium imaging to decode the behavior. Moreover, the analysis of trained encoder-decoder models elicits how the brain processes natural movement. Our approach serves as an example of applying explainable artificial intelligence (Barredo Arrieta et al., 2020) in neuroscience of which importance has been increased for advanced neuroengineering applications and understanding how the brain works (Fellous et al., 2019; Kiani et al., 2022; Lombardi et al., 2021).

### Broader applications to Brain-Machine Interfaces

#### Recovery of movement information from low sampling rate imaging data

Non-invasive or minimally-invasive methods for decoding behaviorally-relevant variables are attractive for the long-term use and stability of BMIs and have recently received massive investment of resources from industry (e.g. Défossez et al., 2023; Ho et al., 2022). However, non-invasive neural recording methods such as electroencephalogram (EEG) and magnetoencephalogram (MEG) primarily reflect strong, synchronous input into a cortical area, lacking fine spatial resolution (Lopes da Silva, 2013). Imaging-based systems can provide finer spatial resolution but often have lower temporal sampling rates (e.g. Tang et al., 2023). Low sampling rates of neural activity are particularly deleterious for real-time neural decoding performance, which is crucial for BMIs. Neural network models can be built to address the low temporal sampling rate by incorporating language models (Tang et al., 2023) or by restructuring data to get sub-frame time resolution (Zhu et al., 2022). Here, we show that population-level neural spiking information is sufficient to reconstruct movement sequences using an encoder-decoder model with longer output than input length. Although here we made use of 2p imaging data, this architecture was originally shown to work on electrophysiological recordings for decoding speech production (Makin et al., 2020), thus transcending a specific recording modality.

#### A small fraction of the neural population is required for decoding

In addition to making use of low-sampling rate data to predict detailed movement trajectories, the encoder-decoder model is an attractive structure for BMI decoding due to the small fraction of the neural population that is relied upon for accurate decoding. This feature, a reliance on approximately ten percent of the recorded population (Figure 4B), can be interpreted as both a strength and a weakness. From a technical perspective, the computational demands for real-time image processing can be reduced once important neurons are identified, as imaging can be focused on gathering just the sparse relevant data. However, over-reliance on a sparse subset of neurons can lead to instability in neural decoding if signals degrade over time. To combat this possibility, the neurons involved in the decode could be periodically re-evaluated and new neurons selected to replace lost or degraded neurons, as has proved successful for electrophysiology-based decoders (Orsborn et al., 2014; Shenoy and Carmena, 2014). Given the large imaging frames we used (covering approximately 1.9 x 1.2 mm of cortex), ten percent of recorded neurons translates into approximately 300 neurons. This is similar to previous work to decode limb trajectories from 2p imaging data (Zhu et al., 2022) and on the high end of the range used to decode electrophysiological data, which often is in the range of 100-300 neurons or recording channels (depending on application; Gilja et al., 2012; Inoue et al., 2018; Makin et al., 2020; Orsborn et al., 2014; Willett et al., 2023). Therefore, both imaging-based decoders and more invasive recording modalities rely upon a similar fraction of neurons (or channels).

#### Simultaneous decoding of ipsilateral and contralateral limbs

The final important application of our work for BMIs is in the simultaneous decoding of both ipsilateral and contralateral limbs from neural data recorded from a single cortical hemisphere, with no significant reduction in decoding accuracy for ipsilateral relative to contralateral limbs (Figure 2) nor any reduction in accuracy in decoding all limbs simultaneously rather than individual models (Figure 3). Our ability to accurately decode ipsilateral limbs reflects prior work showing that neurons within cortical areas ipsilateral to a controlled limb are activated during movements (Ames and Churchland, 2019b; Bundy and Leuthardt, 2019; Bundy et al., 2018). What was more surprising, and advantageous for BMI decoding, is that single-limb and multi-limb models relied upon similar fractions of neurons, as shown by the distribution of neural importance values for the encoder-decoder model (Figure 4A). While the distribution of neural importance assigned to neurons in the multi-limb model is rightward shifted relative to the single-limb model for mice B and C, the shift is small relative to a naive hypothesis that four times the number of neurons would be needed in the multi-limb model to decode four times the number of limbs relative to the single-limb model. Instead, the single-limb and multi-limb decoding showed similar performance with only a small difference in the fraction of the neural population upon which they relied. These results are particularly important for complex BMIs controlling multiple external effectors or for neural prostheses that aim to restore movement in multiple limbs of the body using functional electrical stimulation (Bouton, 2020).

## 4 ACKNOWLEDGEMENTS

The work completed in this article was supported by NSF HDR grant 2117997 and the Ralph W. and Grace M. Showalter Research Trust. We thank Joseph Makin, Eugenio Culurciello, and Vladimir Loncar for helpful discussions regarding the development and training of the encoder-decoder model.

## 5 METHODS

### Animal experiments

All procedures were approved by the Institutional Animal Care and Use Committee at Purdue University. TRE-GCaMP6s (JAX stock #024742) x CaMKII-tTA (JAX stock #007004) male mice aged 3-6 months were used (n=3). Mice were housed in groups of no more than five mice per cage in a temperature and humidity-controlled room on a 12-hour light/dark cycle with *ad libitum* access to food and water. For all surgical procedures, mice were anesthetized with isoflurane at 3% for induction and maintained between 1.5-2.5% to areflexia. All mice were first surgically implanted with a custom titanium headplate (8 mm circular inner diameter) placed over the right cortical hemisphere and secured with dental cement. Following at least three days of recovery, a 5 mm craniectomy over S1 and M1 cortical areas was performed (stereotaxic anatomical coordinates determined from Paxinos & Franklin) and a 5 mm round glass window was placed over the brain and secured with glue. Mice were then given at least three days to recover from surgery before performing 2p imaging.

Mice were given three days to become habituated to the experimenter and experimental room before undergoing habituation to head-fixation while running on an air-lifted Styrofoam ball (Figure 1A). Head-fixation sessions lasted 30 minutes per day for three days. Limb movement during running on the ball was captured with video cameras (The Imaging Source) on the left and right sides of the mouse at 30 frames per second while neural signals were simultaneously recorded with a 2 photon mesoscope (Neurolabware). A Coherent Chameleon Vision Ti:Sapphire laser (920 nm) was used to capture calcium fluorescence of GCaMP6s in layer 2/3 neurons of M1 and S1 (200-300 *µm* below brain surface). Imaging was performed at 1x magnification (approximately 1.9 x 1.2 mm FOV) at 7.81 Hz.

### Data processing

#### Extracting fluorescence traces from two-photon calcium images

Neural signal processing and motion correction was performed with Suite2p (Pachitariu et al., 2016). Suite2p registers neural frames to a reference image, detects cells within a region of interest, extracts fluorescence traces (*F*) from each cell, and performs spike deconvolution.

#### Extracting limb coordinates from behavior video frames

Limb movement was analyzed using DeepLab-Cut (Mathis et al., 2018). DeepLabCut is a markerless pose estimation software that can be used to extract x- and y-coordinates of mouse limbs during running from video frames. Front and hind limbs were manually labeled in 400 video frames, and training ran for 200,000 iterations. The x- and y-coordinates of each limb, *C*^*coord*^ (coord: IF*x*, IF*y*, IH*x*, IH*y*, CF*x*, CF*y*, CH*x*, CH*y*), in each video frame as determined by DeepLabCut were used for behavioral analysis.

#### Data preprocessing for training encoder-decoder models

Fluorescence traces from 2p calcium imaging recordings and limb coordinates from behavior videos were preprocessed for training the encoder-decoder models. Since there were periods where mice ran or remained still, only running periods were used for training and testing. Running periods were defined as periods longer than 30 frames where the difference between the coordinates from the previous frame is larger than 10 pixels. The fluorescence traces were standardized, i.e. *FZ* = (*F − µ*)*/σ* (*µ*: the mean, *σ*: the standard deviation) was used to train the encoder-decoder models. Limb coordinates were normalized (min-max scaling) separately, i.e. 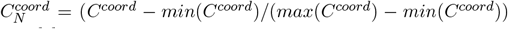 was used to train the encoder-decoder models.

### Applying deep learning

*The network — encoder-decoder model:* To address the low sampling rate of calcium imaging relative to the high speed of natural behavior, we designed a recurrent encoder-decoder network with LSTM (Makin et al., 2020; Sutskever et al., 2014) in which the output sequence length of the decoder is longer than the input sequence length of the encoder (Figure 1F). The encoder consists of LSTM which takes fluorescence traces of neurons extracted from 2p calcium imaging recordings in five neural frames, and the hidden state and the cell state initialized with zeros as inputs. The number of input features was set to the total number of neurons while the number of hidden units and the number of layers were set to hyperparameters to be tuned. The hidden state and the cell state from the final time step of the encoder LSTM are input into the decoder. The decoder consists of LSTM followed by a fully connected layer. The first time step of the decoder LSTM takes a vector with the size of the number of hidden units initialized with zeros, and the hidden state and the cell state from the encoder as inputs. The following time steps get the previous output, the previous hidden state, and the previous cell state as inputs. The number of input features was set to the total number of neurons while the number of hidden units and the number of layers were set to hyperparameters to be tuned. The outputs from the decoder LSTM are fed to the final fully connected layer which converts the input into the vector with a length of two for single-limb decoding models or eight for multi-limb decoding models.

#### Training and hyperparameter tuning

For each animal, a multi-limb decoding model and four single-limb decoding models for each limb (the ipsilateral (right) front limb, the ipsilateral hind limb, the contralateral (left) front limb, and the contralateral hind limb) were trained. The multi-limb decoding model outputs all eight sequences including x- and y-coordinates of all four limbs, and the single-limb decoding model outputs the x-and y-coordinates of one limb. The encoder-decoder models were trained with the first 80% of the whole sequential data. The remaining 20% was used for testing. All networks were trained and tested using PyTorch framework in Python for 30 epochs with early stopping when coefficient of determination (*R*^2^) did not improve more than 0.01. The mean squared error (squared L2 norm) was set as a loss function to minimize by the optimization using Adam.

An additional scheme was required for batch training considering the property of the dataset. Batches were constructed with randomly selected sequences of five neural frames and temporally corresponding limb coordinate sequences. Because of the discrepancy between sampling rates of neural frames and behavior frames, one neural frame matches to various numbers of behavior frames (ranging from two to four frames). Therefore, zero-padding was used to stack limb coordinates with different lengths into batches. A small number (10^*−*6^) was added to zeros in limb coordinates to discriminate between padded zeros. When the output limb coordinates were combined together into one sequence, the padded zeros were removed. The scheme was applied to both training and testing. Overlapped sequences were used for training while non-overlapped sequences were used for testing.

For hyperparameter tuning of each network, tree-structured Parzen Estimator algorithm was used with Optuna (Akiba et al., 2019), a hyperparameter optimization framework. Hyperparameters to be optimized include batch size, learning rate, the number of hidden units of the LSTM layers, and the number of layers of the LSTM layers. For each trial of optimization, the batch size, the number of hidden units, and the number of layers were set as integers randomly sampled between four to 256, five to 1024, and one to three, respectively. The learning rate was set as a float randomly sampled between 10^*−*5^ and 10^*−*1^. The sets of hyperparameters that yielded the best *R*^2^ among 200 trials were selected. Specifications of all networks are shown in Supplemental Table 1.

### Evaluating decoding performance

The decoding performance of the trained encoder-decoder models was evaluated both qualitatively and quantitatively. For the qualitative evaluation, ground truth (*yi, i*: 1, …, *N, N* : the sequence length of the test data) and predictions (*ŷi*) from the encoder-decoder models were visually compared. For the quantitative evaluation, RMSEp was used as a metric. RMSEp is defined as below.

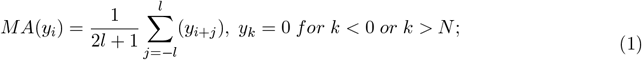

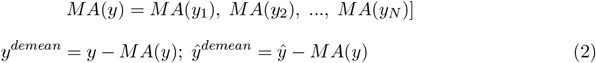

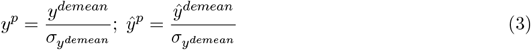

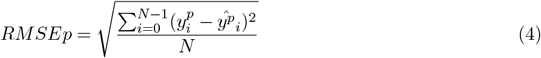

RMSEp was designed considering the high variance in means and scales in the distributions of the different limb coordinates. First, a moving average (MA) was calculated with a window length of 50 (*l* = 25) (equation (1)). Ground truth and predictions were demeaned (*y*^*demean*^ and *ŷ*^*demean*^, respectively) by subtracting moving average of ground truth (*MA*(*y*)) (equation (2)). Processed ground truth (*y*^*p*^) and processed predictions (*ŷ*^*p*^) were obtained by dividing ground truth and predictions with the standard deviation of *y*^*demean*^ 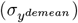 (equation (3)). RMSEp was obtained by calculating root mean squared errors using the processed data (*y*^*p*^ and *ŷ*^*p*^) (equation (4)). Normal root mean squared error is not an appropriate measure for comparison between limbs or subjects because each sequence of limb coordinates have different value ranges. While *R*^2^ can be considered to show general regression performance, it might not be appropriate for non-linear data (Spiess and Neumeyer, 2010) such as limb coordinates in this study. The two-sided independent two-sample t-test was performed to check the statistical significance between different cases.

### Relative neural importance

The neural importance was obtained by calculating a norm of the gradients of the loss function with respect to each neuron across the sequences of the test data (Makin et al., 2020; Simonyan et al., 2014). The calculated norm of the gradients was min-max scaled between zero and one to represent the relative importance. The idea is that the gradient shows how much the loss function changes by small changes in signals from each neuron, which ultimately shows how each neuron is important in the decoding process of the trained encoder-decoder models (Makin et al., 2020). The distributions of importance from multi-limb and single-limb (combined across all limbs and each limb) decoding models were shown in histograms and compared. The two-sample Kolmogorov-Smirnov test was performed to test the significance of differences between the distributions. The spatial dispersion or sparsity of the important neurons was evaluated by averaging the relative importance over each grid with a length of 50 pixels on the imaging plane.

Furthermore, the effect of the importance of neurons in decoding was evaluated by measuring the decoding performance of subsets of neurons (10%, 25%, 50%, 75%, and 90% of the total) that were most important, least important, and sampled randomly. For each subset, the most and the least important neurons were sampled and 51 subsets of neurons were randomly sampled, of which one was used for the hyperparameter optimization. With the fixed set of hyperparameters, a total of 52 multi-limb models were trained separately with each subset of neurons. A two-sided independent two-sample t-test was performed to check the statistical significance between the adjacent number cases.

### Neural tuning to limb position

For each neuron, we generated a *tuning curve* that described the average response of that neuron to the positions of each of the four limbs. In these analysis, neural responses were taken to be the normalized, deconvolved fluorescence traces used to train the encoder-decoder model. Since our recordings of neural activity were at a lower frequency than recordings of limb movements, for this analysis we repeated the value of neural activity for the two to four behavioral frames that matched up with each neural imaging frame. To calculate tuning curves from neural activity, we first divided the position of each limb into eleven equally sized bins (separately for x and y position), ranging from *µ −* 2*σ* to *µ* + 2*σ* of the normalized, de-meaned trajectory for each limb. Next, we calculated the average activity of that neuron whenever that limb was in that coordinate range (i.e., that bin). From the tuning curves for each limb, we defined a neuron’s *preferred limb* as the limb to which the neuron had the largest response across all eight tuning curves. Within the tuning curve for a neuron’s preferred limb, we further defined a neuron’s *preferred position* as the coordinates to which the neuron had the largest response. The number of peaks within a tuning curve was found as the number of points within the curve that were higher than two surrounding points (using the MATLAB *findpeaks* function) and was calculated for neurons that had a tuning height of at least 0.3. Given that tuning was calculated using normalized neural activity, a tuning height of 0.3 means that the difference between the minimum and maximum average response to limb position is at least a difference of 0.3 standard deviations of that neuron’s activity level.

For each tuning curve, we calculated two measures of tuning: tuning height and circular variance. *Tuning height* was defined as the difference between the maximum and minimum of each tuning curve and thus served to measure the difference between the strongest and weakest response of the neuron. *Circular variance* is a form of directional statistics that assumes responses are wrapped around a circle (Fisher, 1993). We adopt this terminology because the oscillatory limb movements of the mouse can be thought of as movement around a circle. Extending this analogy, we can describe the most anterior or ventral limb coordinates at 0 degrees and the most posterior or dorsal limb coordinates at 180 degrees. Circular variance measures the spread of a distribution around a circle and can be calculated as in Equation 5. A large circular variance corresponds to larger spread of the tuning curve while a small circular variance corresponds to narrower tuning.

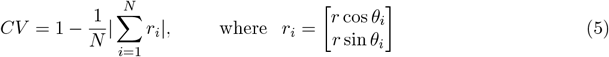

### Signal processing of limb movements and predictions

A power spectrum analysis of true and predicted limb movements was conducted in MATLAB using the *fft* function. A separate analysis was computed for each limb in both x- and y-coordinates. Given that limb movements were sampled at 30 Hz, the computed power spectrum went between 1 Hz and 15 Hz — half the behavioral sampling frequency. Finally, to determine which components of the limb movements are best captured by the encoder-decoder model, we computed Pearson correlation coefficient between the RMSEp values and the power within each frequency band across limbs, generating fifteen correlation coefficients for each of the fifteen frequency components that were analyzed. Correlations were computed separately for each mouse.

## Notes

### Competing Interest Statement

The authors have declared no competing interest.

