## Supplemental materials for "Decoding multi-limb movements from low temporal resolution calcium imaging using deep learning"

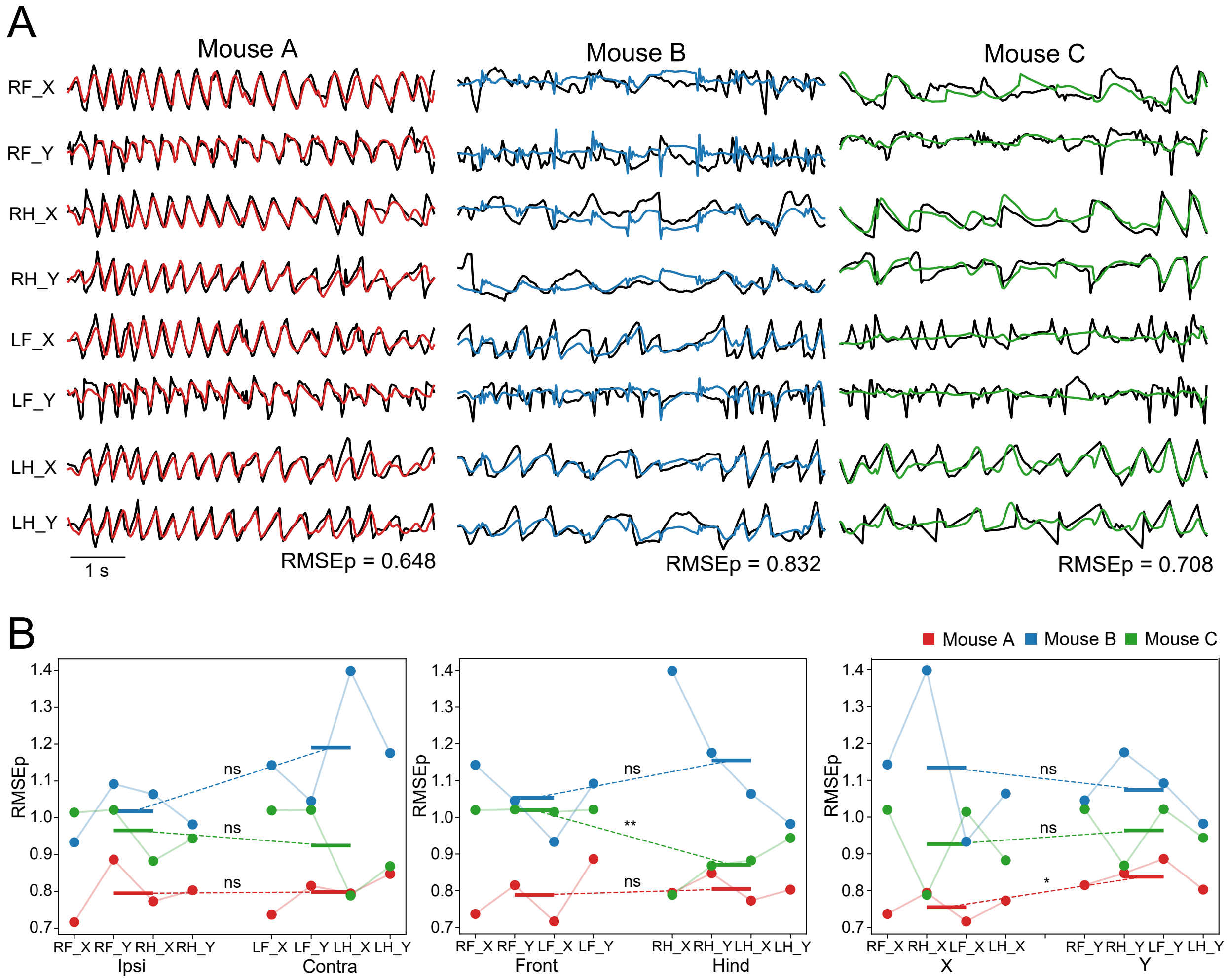


Supplemental Figure 1: Multi-limb decoding of contralateral vs. ipsilateral limbs. (A) Sample traces showing ground truth (black) and model predictions (colored) for contralateral (right) and ipsilateral (left) movement in mice A, B, and C. (B) Limb prediction RMSEp across all mice and all limbs, grouped by ipsilateral vs. contralateral (left) ($p = 0.505$; Mouse A: $p = 0.933$; Mouse B: $p = 0.083$; Mouse C: $p = 0.560$; t-test), front vs. hind limbs (middle) ($p = 0.880$; Mouse A: $p = 0.718$; Mouse B: $p = 0.352$; Mouse C: $p = 0.004$; t-test), and x-axis vs. y-axis (right) ($p = 0.772$; Mouse A: $p = 0.018$; Mouse B: $p = 0.587$; Mouse C: $p = 0.596$; t-test).

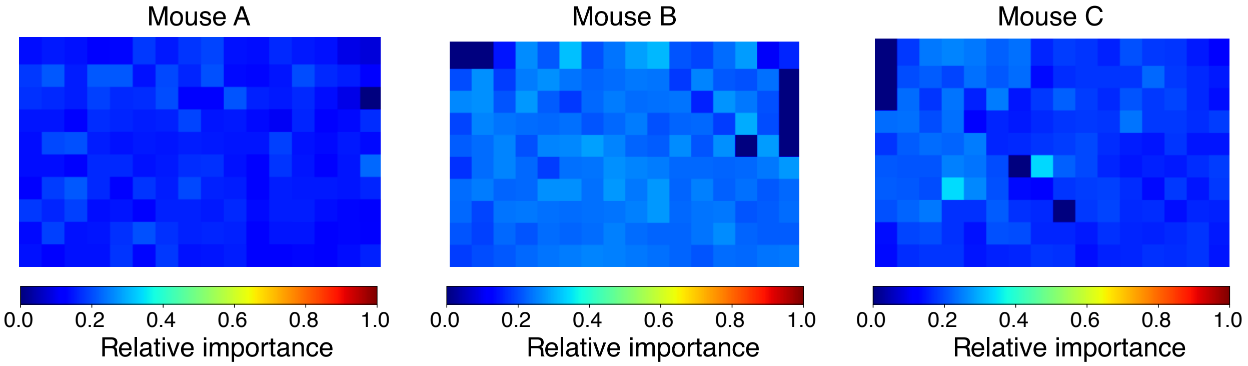


Supplemental Figure 2: Normalized neural importance spatially averaged over 116 by 116 $\mu$m bins.

Supplemental Table 1: Specifications of trained networks. Input: the number of input features; HU: the number of hidden units; Layer: the number of LSTM layers; Output: the output size of the last fully connected layer; LR: learning rate

| Network | Input | HU | Layer | Output | LR | Batch size |
|---|---|---|---|---|---|---|
| Mouse A, single-limb, IF | 3327 | 829 | 3 | 2 | 0.001957 | 12 |
| Mouse A, single-limb, IH | 3327 | 797 | 3 | 2 | 0.002278 | 40 |
| Mouse A, single-limb, CF | 3327 | 458 | 2 | 2 | 0.001907 | 28 |
| Mouse A, single-limb, CH | 3327 | 917 | 2 | 2 | 0.001053 | 16 |
| Mouse A, multi-limb | 3327 | 795 | 2 | 2 | 0.001930 | 12 |
| Mouse A, multi-limb, 10% | 332 | 1019 | 2 | 8 | 0.001035 | 104 |
| Mouse A, multi-limb, 25% | 831 | 991 | 2 | 8 | 0.000349 | 4 |
| Mouse A, multi-limb, 50% | 1663 | 437 | 3 | 8 | 0.001615 | 20 |
| Mouse A, multi-limb, 75% | 2495 | 687 | 2 | 8 | 0.001229 | 12 |
| Mouse A, multi-limb, 90% | 2994 | 698 | 3 | 8 | 0.001672 | 32 |
| Mouse B, single-limb, IF | 2978 | 590 | 3 | 2 | 0.000394 | 32 |
| Mouse B, single-limb, IH | 2978 | 539 | 3 | 2 | 0.000216 | 8 |
| Mouse B, single-limb, CF | 2978 | 394 | 2 | 2 | 0.003157 | 12 |
| Mouse B, single-limb, CH | 2978 | 632 | 3 | 2 | 0.000630 | 4 |
| Mouse B, multi-limb | 2978 | 628 | 1 | 8 | 0.001522 | 12 |
| Mouse B, multi-limb, 10% | 297 | 256 | 3 | 8 | 0.003771 | 8 |
| Mouse B, multi-limb, 25% | 744 | 838 | 2 | 8 | 0.000246 | 4 |
| Mouse B, multi-limb, 50% | 1489 | 517 | 2 | 8 | 0.001808 | 96 |
| Mouse B, multi-limb, 75% | 2233 | 845 | 2 | 8 | 0.000412 | 4 |
| Mouse B, multi-limb, 90% | 2680 | 544 | 3 | 8 | 0.000166 | 4 |
| Mouse C, single-limb, IF | 3314 | 914 | 3 | 2 | 0.001760 | 24 |
| Mouse C, single-limb, IH | 3314 | 992 | 1 | 2 | 0.001404 | 12 |
| Mouse C, single-limb, CF | 3314 | 970 | 3 | 2 | 0.000211 | 8 |
| Mouse C, single-limb, CH | 3314 | 900 | 2 | 2 | 0.000161 | 8 |
| Mouse C, multi-limb | 3314 | 397 | 3 | 8 | 0.002211 | 28 |
| Mouse C, multi-limb, 10% | 331 | 382 | 3 | 8 | 0.002224 | 16 |
| Mouse C, multi-limb, 25% | 828 | 783 | 3 | 8 | 0.001640 | 76 |
| Mouse C, multi-limb, 50% | 1657 | 247 | 3 | 8 | 0.001978 | 24 |
| Mouse C, multi-limb, 75% | 2485 | 467 | 2 | 8 | 0.001742 | 20 |
| Mouse C, multi-limb, 90% | 2982 | 923 | 3 | 8 | 0.000194 | 32 |
